# Galangin and Caffeic acid inhibit Methylglyoxal-induced Advanced Glycation End Product formation in Bovine Serum Albumin

**DOI:** 10.64898/2026.07.09.737425

**Authors:** Nupur Kanojia, A.B. Tiku

**Affiliations:** Radiation and Therapeutics Laboratory, School of Life Sciences, Jawaharlal Nehru University, New Delhi-110067, India

**Keywords:** Advanced Glycation End Products (AGEs), Glycation, Methylglyoxal, Aggregation, Phytochemicals, Galangin, Caffeic Acid

## Abstract

Glycation, a non-enzymatic reaction occurring between sugars and biological macromolecules, plays a critical role in ageing and disease pathogenesis. Methylglyoxal (MG) is a highly reactive α-oxoaldehyde that leads to the formation of endogenous advanced glycation end products (AGEs). These AGEs are associated with diabetes and many other diseases, including neurodegeneration and cancer. This is often through interactions with the receptor for advanced glycation end products (RAGE). Inhibition of glycation/AGEs formation using natural products to target cancer is an area of recent interest.

In vitro AGEs formation was observed by browning of samples, increased fluorescence, and carbonyl stress. MG induced changes in the structure of BSA were analysed using electrophoresis, spectroscopy, TEM, AFM, DLS, and CD spectroscopy. Our results show that AGEs form random structures, oligomeric aggregates, and β-sheets. Thioflavin T and Congo red staining further validated these findings. Galangin and Caffeic acid demonstrated significant antiglycation activity, suppressing AGEs formation *in vitro*.

**Graphical Abstract:** 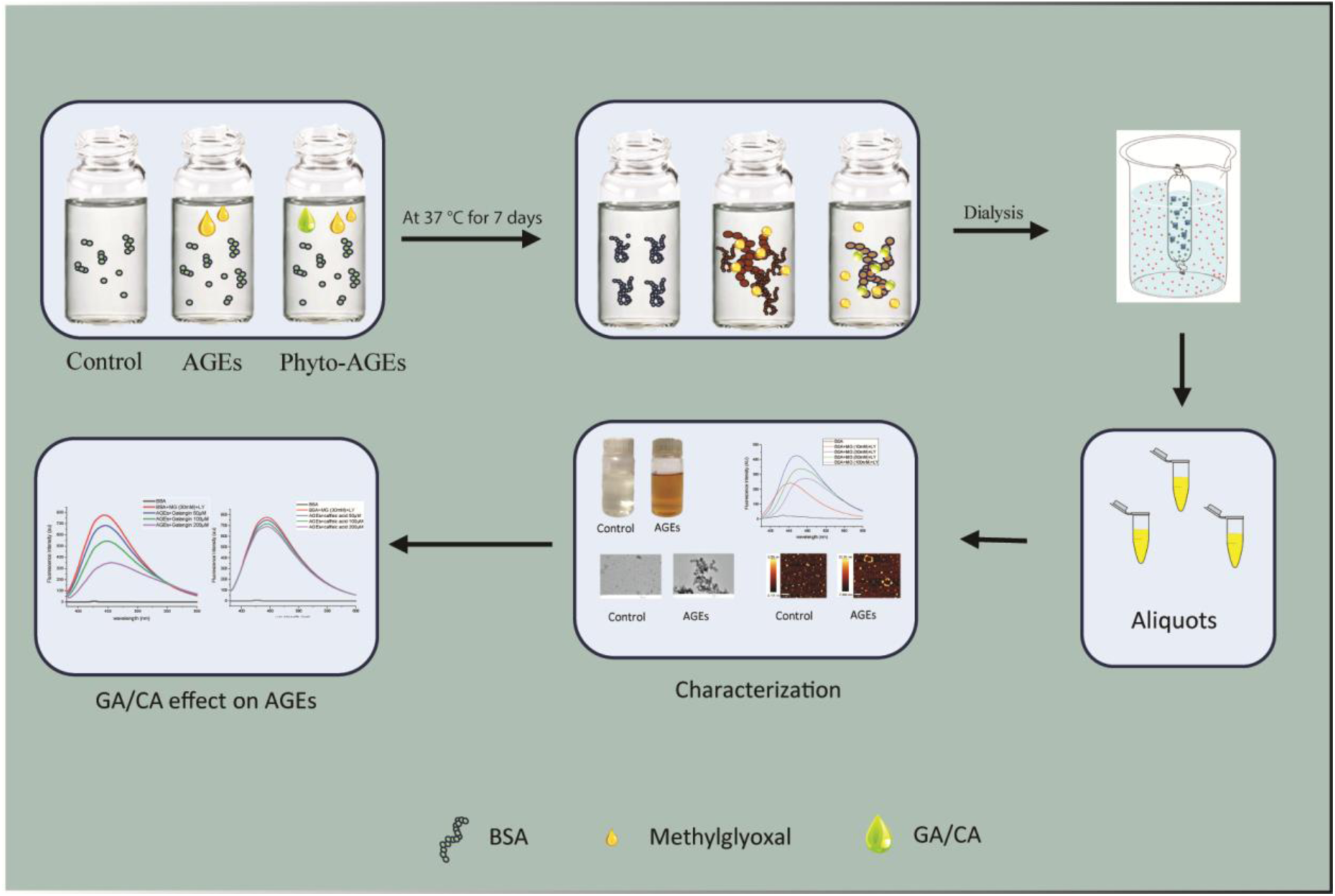

**Highlights:**

- Methylglyoxal-induced Advanced Glycation End Products were prepared in vitro
- Methylglyoxal -induced structural modifications in BSA
- AGEs were characterised using various parameters
- Both fluorescent and non-fluorescent AGEs were formed.
- Phytochemical treatment induced inhibition of AGEs formation

## 1. Introduction

Glycation results from the spontaneous, non-enzymatic reaction of reducing sugars and macromolecules. The sugars, primarily glucose, react with the N-terminal of various amino acids and give rise to an initial unstable Schiff’s base, followed by rearrangements leading to the generation of an irreversible adduct known as advanced glycation end product (AGEs).

Several external factors are also responsible for the AGEs pool in the body, which include smoking, high carbohydrate diet, grilled food, high-temperature cooking, and a sedentary lifestyle. Methylglyoxal (MG), an endogenous toxic offshoot of glycolysis, is also responsible for the formation of AGEs. MG, glyoxal, and 3-deoxyglucosone are highly reactive α-oxoaldehyde metabolites that generate AGEs [1]. MG has approximately 20,000 more reactivity towards macromolecules, compared to glucose [2].

The MG-induced modifications leading to the formation of AGEs are irreversible and result in protein inactivation and dysfunction. Various AGEs present in the physiological system are CML, CEL [Ne-(1-carboxyethyl)-lysine], pyrraline, MOLD [methylglyoxal-derived lysine dimer, 1,3-di(Ne-lysino)-4-methyl-imidazolium salt], GOLD [glyoxal-derived lysine dimer, 1,3-di(Ne-lysino)imidazolium salt] and DOLD [3-deoxygluco-sone-derived lysine dimer, 1,3-di(Ne-lysino)-4-(2,3,4-trihydroxybutyl)-imidazolium salt], THP [arginine-derived tetra hydro pyrimidine], pentosidine, and argpyrimidine. MGdG is formed via interaction of MG with DNA. Adducts formed can result in DNA damage, mutations, DNA-DNA crosslinks, and DNA-protein crosslinks. Approximately 9 adducts per 106 nucleotides are produced *in vivo,* and their increased occurrence is often associated with various diseases [3].

In normal physiological conditions, elimination of AGEs occurs via the kidneys to maintain a steady state. However, ageing and other extrinsic factors lead to excessive accumulation of AGEs, resulting in oxidation, inflammation, and various diseases, including cancer. AGEs promote oxidative damage and stress, which activate transcription factors, causing increased production of proinflammatory and inflammatory mediators. Recently, their relationship with receptors for advanced glycation end products (RAGE) was found to be involved in the initiation and progression of cancer [4–8]

Various antiglycation agents have been explored to overcome the burden of glycation-induced AGEs. Synthetic antiglycation agents like aminoguanidine, inositol, diclofenac, buformin, pyridoxamine, alagebrium, and metformin have been shown to inhibit the Maillard reaction, and limit the development of intermediates of glycation [9]. However, considering the complications linked with synthetic molecules in clinical trials, natural products have been explored for their AGE-inhibiting properties. Natural compounds have been found to be safe and non-toxic in the treatment of glycation-induced diseases [10]. Many phytochemicals, besides having antioxidant activity, have also been reported to have antiglycation properties. They have been reported to inhibit AGEs intermediates like carbonyl compounds, Schiff’s base, and amadori compounds, ultimately leading to blockage of AGE-RAGE interaction [8]. Dietary polyphenols like Ellagic acid inhibit AGEs (fluorescent and non-fluorescent) by acting as carbonyl scavengers. Moreover, ellagic acid prevents the formation of beta-sheets in haemoglobin and lysozyme [11]. Resveratrol traps MG and acts as a free radical scavenger. Additionally, ellagic acid, kaempferol, epicatechin, and catechin not only prevent AGEs formation but also bind effectively with RAGE and prevent AGEs-RAGE induced downstream signalling pathways [12]. Curcumin, an active principle of turmeric, was found to lower the blood glucose levels in mice with type 2 diabetes [13]. Other flavonoids like gallic acid, genistein, etc, inhibit AGEs formation by MG entrapping. They are also glycoxidation inhibitors [14].

Galangin (GA) a phytochemical present in propolis and roots of *Alpinia officinarum*, has been reported to have significant antihyperlipidemic, antidiabetic, anti-inflammatory, antioxidant, antiviral, antimicrobial, and antifungal properties [15–17]. Caffeic Acid (CA) another well-known phenolic compound found in almost all species of plants like olives, berries, carrots, potatoes, coffee, red wine, apple cider vinegar, tea, coffee, and propolis has anti-inflammatory, antioxidant, anti-microbial, and anti-neoplastic properties [18]. CA is reported to show antihyperglycemic effects in streptozotocin-induced diabetic rats [19]. Apart from having many biological activities, both these phytochemicals are reported to possess anti diabetic properties [9–11]. However, mechanism of the antiglycation effects of these phytochemicals is not clear.

Therefore in the present study we have evaluated GA and CA for their antiglycation properties using the methylglyoxal-BSA system [15]. Targeting AGEs formation is a promising therapeutic intervention in various ageing and AGEs-related diseases, including cancer.

## 2. Materials and Methods

### 2.1. Material

All materials, BSA (Hi-Media), Methylglyoxal, Sodium azide, Lysine, GA, CA, Trichloro Acetic Acid (TCA), 2,4-Dinitrophenylhydrazine (2,4-DNP), Guanidine Hydrochloride, Anti-AGEs antibody (ab23722), Thioflavin T stain, and Congo-Red stain, were purchased from Sigma-Aldrich, unless otherwise mentioned

### 2.2. Methodology

#### 2.2.1. Preparation of MG-BSA (AGEs)

AGEs were prepared as per Ahmed et al [20] with slight modifications. To generate AGEs in vitro, a mixture containing 2 mg/mL of BSA, 3 mM sodium azide, 1.5 mM lysine, and varying concentrations of MG (10, 30, 50, and 100 mM) was prepared using 100 mM sodium phosphate buffer (pH 7.4). The MG-BSA samples were transferred into glass vials and incubated at 37 °C for seven days. As a control (sham BSA), BSA alone was incubated under the same conditions without MG and lysine. After incubation, all samples underwent thorough dialysis against sodium phosphate buffer for 72 hours. The Bradford assay was used to determine protein concentration.

#### 2.2.2. Phytochemical treatment

GA and CA were dissolved in DMSO (final conc. 0.1%) and added to BSA before adding MG. Rest all the experimental conditions were similar to BSA and AGEs control. The Bradford assay was used to determine protein concentration.

#### 2.2.3. Fluorescence Spectroscopy

The fluorescence spectrum of MG-BSA was observed at Ex_370nm_ using a Shimadzu fluorescence spectrophotometer, RF-5300, Japan, after digesting the samples with 6N HCl [20]. The sample concentration was 1mg/mL. Fresh BSA and Sham BSA were used as controls. Emission spectra were recorded at excitation 325 nm, 485 nm, and 285 nm for argpyrimidine, pentosidine, and tryptophan quenching [21].

#### 2.2.4. UV-Vis Spectroscopy

The UV-Vis spectrum of MG-BSA (AGEs) of varying concentrations and control BSA was monitored using a Shimadzu 1800 spectrophotometer. Sodium phosphate buffer 0.1M at pH 7.4 was used as blank with a 1cm pathlength. The total volume was maintained at 1 mL.

#### 2.2.5. Di-Carbonyl detection

The carbonyl content of AGEs of varying MG concentrations and controls was estimated using a standard protocol [22]. In brief, 1mg/mL of protein solution was taken and precipitated with freshly prepared TCA (20%) and spun down at 2655g for 5 minutes. The pellet was resuspended in 10 mM 2,4-DNP solution in 2M HCl and kept in the dark for 1 hour. The samples were vortexed every 10 minutes. Samples were again centrifuged at 2655g, and the pellet was washed with 1:1 (v/v) mixture of ethyl acetate and ethanol. 6M guanidine hydrochloride was added to each vial, and samples were kept for 15 minutes at 37 °C. Absorbance was taken at 366nm using 6M guanidine hydrochloride solution as a blank using Shimadzu 1800 UV-Vis spectrophotometer, Japan.

#### 2.2.6. SDS-PAGE (Gel electrophoresis)

SDS-PAGE was performed according to Laemmli’s protocol with minor alterations [23]. In brief, 10µg of sample was loaded and the gel was run at 80V using a mini-PROTEIN II Bio-Rad electrophoresis system till the samples were completely resolved. Gel was stained with Coomassie Brilliant Blue (CBB) for 15-20 minutes with continuous rocking at room temperature and later destained till the bands were properly visible.

#### 2.2.7. Native Gel Electrophoresis

For the aggregates, native polyacrylamide gel electrophoresis was carried out using a 10% acrylamide gel in a Bio-Rad mini-PROTEIN II system. The electrophoresis was performed at a consistent voltage between 60 and 70V. The gel was stained with CBB dye and destained till the bands were visible [24].

#### 2.2.8. Western Blotting

Western blotting was done to verify the *in vitro* formation of MG-BSA (AGEs). In brief, proteins were denatured by heating at 95 °C for 5 minutes with sample buffer . The protein samples were then resolved using 10% SDS-PAGE at 80V for 2–3 hours and transferred using a PVDF membrane. 5% BSA was used as a blocking buffer. The blot was incubated with primary antibody (ab23722) diluted 1:1000 overnight at 4 °C, followed by three washings with TBST. The membrane was incubated with a secondary antibody, followed by washing. Signal detection was carried out using enhanced chemiluminescence and visualised with the Chemiluminescent Imager (G Box Chemi XRQ), Syngene, UK.

#### 2.2.9. Dynamic Light Scattering Spectroscopy

The hydrodynamic size was characterized using Malvern Nano-ZS90 Zetasizer, Malvern Instruments Ltd., Worcestershire, UK, in 12 mm square disposable cuvette. Samples were diluted using filtered (0.22µm) buffer. The intensity of laser light scattered at a 90-degree angle was tracked over approximately one hour (each reading of 100 runs) at 25 °C [24].

#### 2.2.10. Circular Dichroism

The BSA and AGEs samples were subjected to Circular dichroism (CD) analysis using an Applied PhotoPhysics Chirascan equipped with a Peltier temperature controller. The sample concentration was kept at 0.75 μg/mL and placed in a CD cuvette. The path length was 2 mm. The buffer used to initiate the reaction served as the reference. All measurements and readings were done at ambient temperature. Each CD spectrum presented represents the average of three separate scans, recorded in the wavelength range of 200–260 nm. CD data were analysed using the BeStSel web server (https://bestsel.elte.hu).

#### 2.2.11. Thioflavin T Staining (Fluorescence Spectroscopy)

4 µM protein sample was incubated with ThT stain (Thioflavin T, concentration, 20 µM). The fluorescence emission spectrum was recorded at excitation/emission wavelengths of λEx 440 nm / λEm 490 nm. All aggregation assays were conducted at room temperature. ThT fluorescence intensity was measured using a Shimadzu RF-5300 fluorescence spectrophotometer (Japan) .

#### 2.2.12. Congo-Red staining (UV-Vis Spectroscopy)

To evaluate amyloid formation in the AGE samples, Congo Red (CR) dye was used. 200µg of samples were stained with 30 µM CR dye and incubated at RT for 30 minutes. Absorbance spectra were then recorded in the 400–600 nm range using a UV-Vis spectrophotometer.

#### 2.2.13. Transmission Electron Microscopy

AGEs were examined under a Transmission electron microscope (TEM JEOL MEM-2100F/ HR Tokyo, Japan). Initially, samples were dried on a copper grid, followed by washing with water and air drying. The air-dried grids coated with BSA and AGEs were stained with 2% (w/v) aqueous uranyl acetate for 10 minutes, followed by gentle washing and drying. The grids were analysed and imaged using an electron microscope operated at an accelerating voltage of 200 kV.

#### 2.2.14. Atomic Force Microscopy (AFM)

For AFM measurements, protein samples were smeared on freshly cleaved mica sheets, and air dried. The AFM topography images were captured under proper working atmosphere. Software PROJECT 4.0 was used for data analysis.

#### 2.2.15. Statistical Analysis

All experiments were conducted a minimum of three times. For statistical analysis, a one-way ANOVA was used to compare the means of different groups. Data are presented as the mean ± SEM from three independent experiments. P-value of <0.05 was considered statistically significant.

## 3. Results and Discussions

### 3.1. Detection of Advanced Glycation End products (AGEs)

Incubation of BSA with methylglyoxal for 7 days at 37 °C resulted in colour change with no visible precipitation, indicating the formation of AGEs. Change in colour is the first characteristic trait that is observed in the formation of AGEs (Figure 1a). The brown colour observed is due to the formation of melanoidins. They are the high molecular weight, amino aggregates generated in the ultimate stage of the Maillard reaction [25]. No significant colour change was observed for samples containing only MG or only lysine. However slight yellowish colour was observed in samples containing both MG and lysine. A dark brown colouration was observed in samples containing MG with BSA, confirming the formation of AGEs. MG is capable of modifying both lysine and arginine residues at high concentration [26].

**Figure 1.**
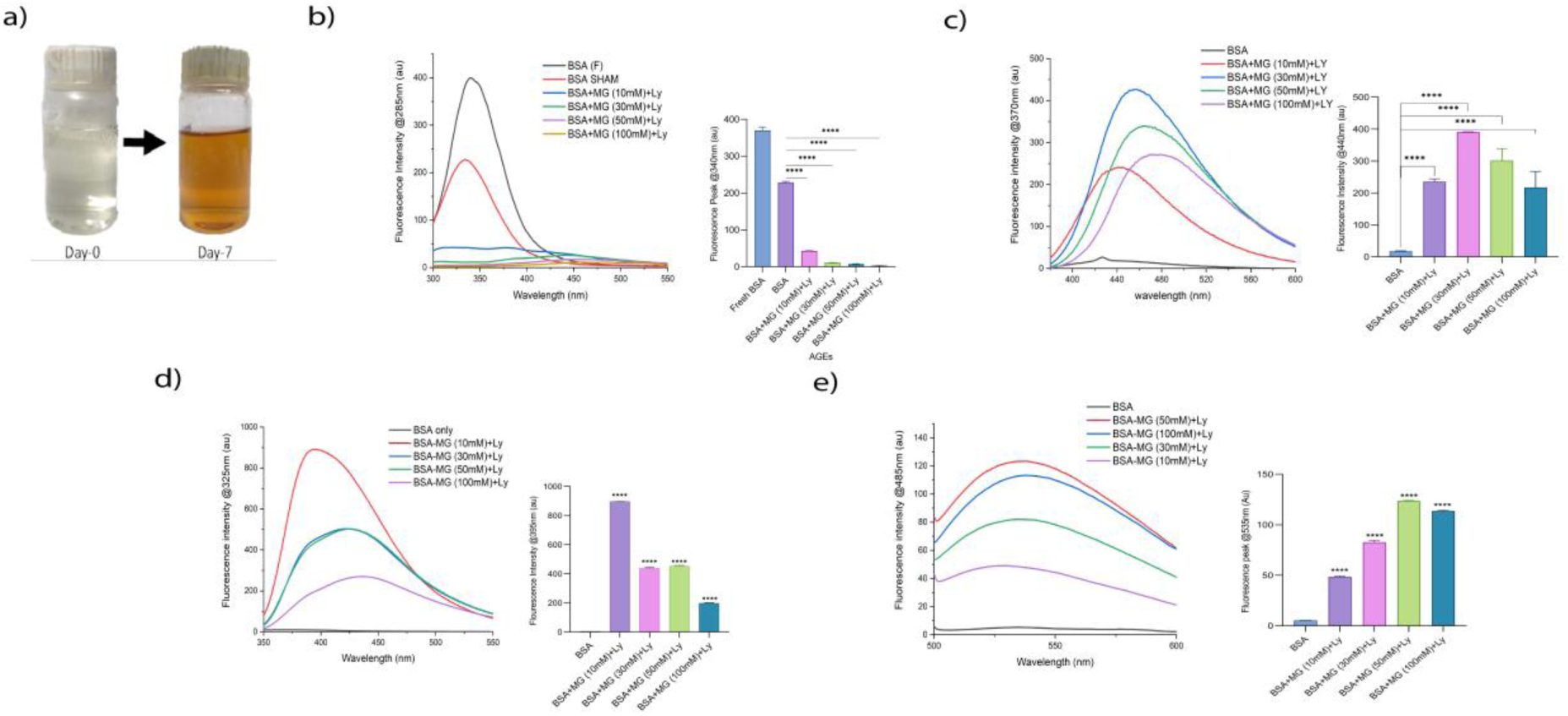
Detection of AGEs, **(a)** Change in colour (colourless to brown). **(b)** Fluorescence Emission Spectrum of BSA vs MG-BSA (AGEs) at various MG concentrations (10mM, 30mM, 50mM, and 100mM). **(c)** AGEs specific fluorescence spectrum at excitation wavelengths 325nm, and **(d)** 485nm, **(e)** The fluorescence quenching spectrum of native BSA and MG-BSA (AGEs), recorded at an excitation wavelength of 285nm, Significance level ****p<0.0001, with respect to control (n=3).

### 3.2. Fluorescence Spectroscopy

AGEs are categorized into two categories based on the presence or absence of fluorescence. Pentosidine and argpyrimidine are fluorescent and CML and CEL are non-fluorescent AGEs. [27]. BSA also has intrinsic fluorescence at a wavelength of λEx285/Em350nm due to the presence of amino acid tryptophan. The fluorescence quenching experiment showed a decrease in fluorescence intensity (Figure 1b). A dose-dependent decrease in intrinsic fluorescence could be due to structural changes in BSA induced by MG [20].The total AGEs formed were observed at λ_Ex370/Em450nm_ using fluorescence spectroscopy. An increase in fluorescence was noted in the AGEs) samples (treated with 6N HCl to release bound AGEs from serum albumin) when compared to BSA alone [28]. The difference in fluorescence between sham control BSA and MG-BSA samples is attributed to the formation of fluorescent AGEs [16]. The maximum fluorescence was observed for 30mM concentration of MG (Figure 1c). The emission fluorescence spectrum at λ_Ex325nm/Em395nm_ suggests the presence of Argpyrimidine, and fluorescence at λ_Ex485nm/Em535nm_ shows the presence of pentosidine AGEs. The presence of both pentosidine and argpyrimidine AGEs was observed in our sample. (Figure 1d,e) [21].

### 3.3. UV-Vis Spectroscopy

Modification of BSA with different concentrations of MG was also detected using UV-Vis spectroscopy. A characteristic peak of BSA was observed at 280 nm, which is due to tyrosine and tryptophan, aromatic amino acids. By distorting natural protein structure, glycation exposes aromatic amino acid residues and alters their microenvironment, which ultimately increases absorbance. An increase in absorbance indicates the formation of AGEs [29]. However, when BSA was incubated with different concentrations of MG, a strong increase in absorbance was observed (Figure 2). Besides, an additional peak at 320–330 nm was also observed. This additional peak may be due to modified arginine residues of BSA [30].

**Figure 2.**
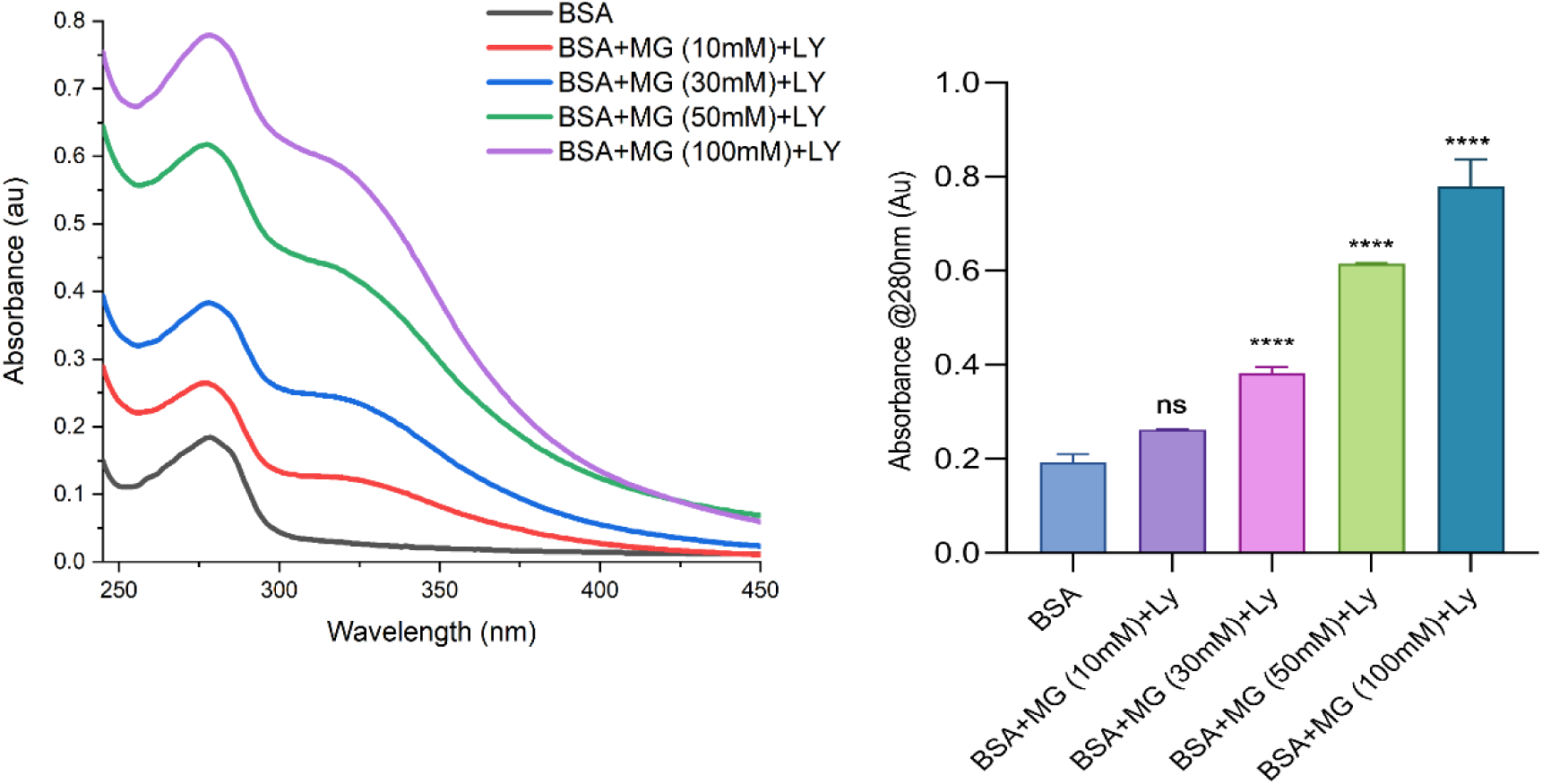
UV-Vis spectrum of BSA (control) and MG-BSA (AGEs) at different MG concentrations (10mM, 30mM, 50mM, and 100mM). The spectrum was recorded in the range of 250 nm-450 nm, marking the additional peak in MG-BSA between 320-330 nm, Significance level ****p<0.0001, with respect to control (n=3).

### 3.4. Carbonyl Content

The carbonyl content is a biomarker for oxidative damage and reactive carbonyl species (RCS). Carbonyl groups are highly reactive and form chemically stable compounds that have a variety of functional consequences. Glycation/Oxidation of side chains of proteins (pro, arg, lys, and Thr) result in the production of these compounds [31]. Thus, elevated carbonyl content is a characteristic of AGEs formation in a physiological system and *vitro* models. In the present study, an increased carbonyl content was also observed in all the samples. Carbonyl content was significantly increased in MG-BSA (AGEs) compared to BSA alone. A 3-fold increase in carbonyl content was observed in 30 mM MG-BSA compared to the BSA control. A dose-dependent increase in carbonyl content was observed only up to 30mM concentration of MG, but no significant difference was found between the total carbonyl content of AGEs prepared with 30mM, 50mM, and 100mM (Figure 3) [30]. The concentration of substrate and glycating agent could be one of the limiting factors as BSA has a strict number of arginine and lysine subunits [32].

**Figure 3.**
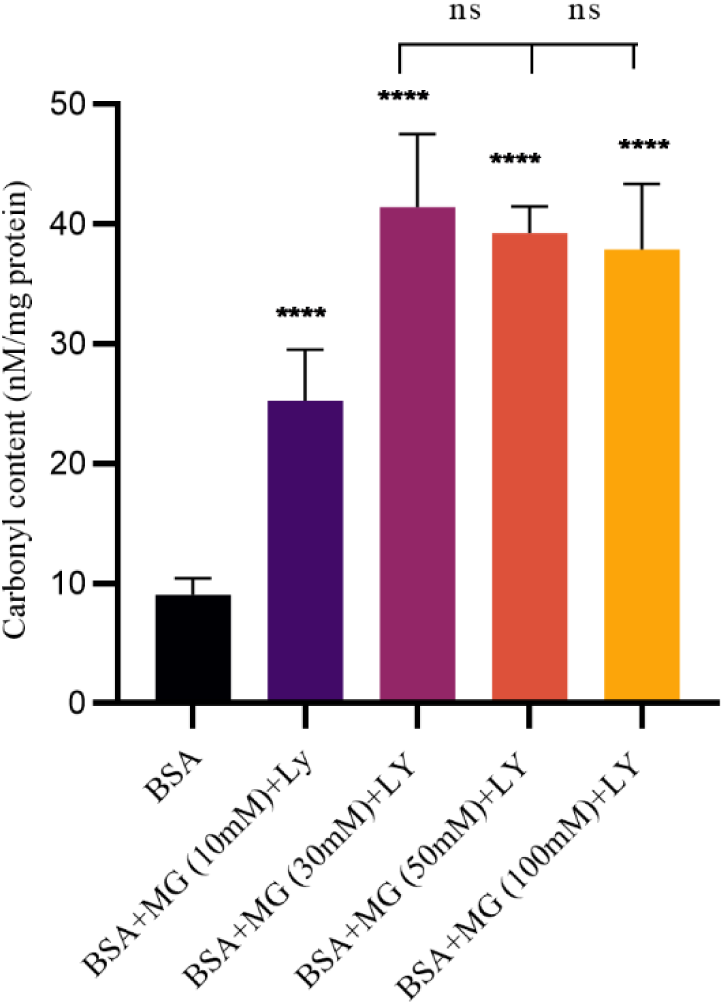
Carbonyl content of BSA (control) and MG-BSA (AGEs) at various MG concentrations (10mM, 30mM, 50mM, and 100mM). Significance level ****p<0.0001, with respect to control (n=3).

### 3.5. Characterization using Electrophoresis

Glycation leads to structural modifications of free amino groups in the proteins. To check the molecular integrity of MG-BSA (AGEs), native PAGE (10%) was run. Untreated BSA showed monomeric and dimeric bands due to covalent and non-covalent aggregation in solution. A significant change in mobility and heterogeneity was observed in MG-BSA (AGEs). An increased electrophoretic mobility and diffused bands of MG-BSA (AGEs) suggested the formation of cross-linking products. These observations indicate considerable modification of the protein with increasing concentration of MG. A smear of the protein band observed at the top of the lane at higher concentration of MG (30mM) suggested formation of larger aggregates. Such changes were only observed in MG-BSA (AGEs) and were absent in control BSA (Figure 4a). Although no visible aggregation or precipitation was observed in the AGEs solution (Figure 1a), bands were observed in native PAGE suggesting that AGEs formed, soluble as well as a mixture of several subtypes of aggregates. The increase in electrophoretic mobility could be due to loss of positive charge(s) as MG modification have been found to be associated with increased electrophoretic mobility of proteins [33–35]. Glycation of haemoglobin by fructose/ glyoxal has been reported to enhance the electrophoretic mobility of the proteins due to loss of positive charge(s) [35]. Further bands of MG-BSA (AGEs) and BSA alone in SDS PAGE are at around 66kD, suggesting that AGEs are stable under reducing conditions and increased negative charge is due to structural changes in BSA. (Figure 4b) [36]. AGEs formation was further confirmed by performing western blotting of MG-BSA (AGEs) against anti-AGEs antibody. Samples were loaded in increasing concentrations from 5µg to 50 µg. The positive binding of antibody confirms AGEs formation (Figure 4c).

**Figure 4.**
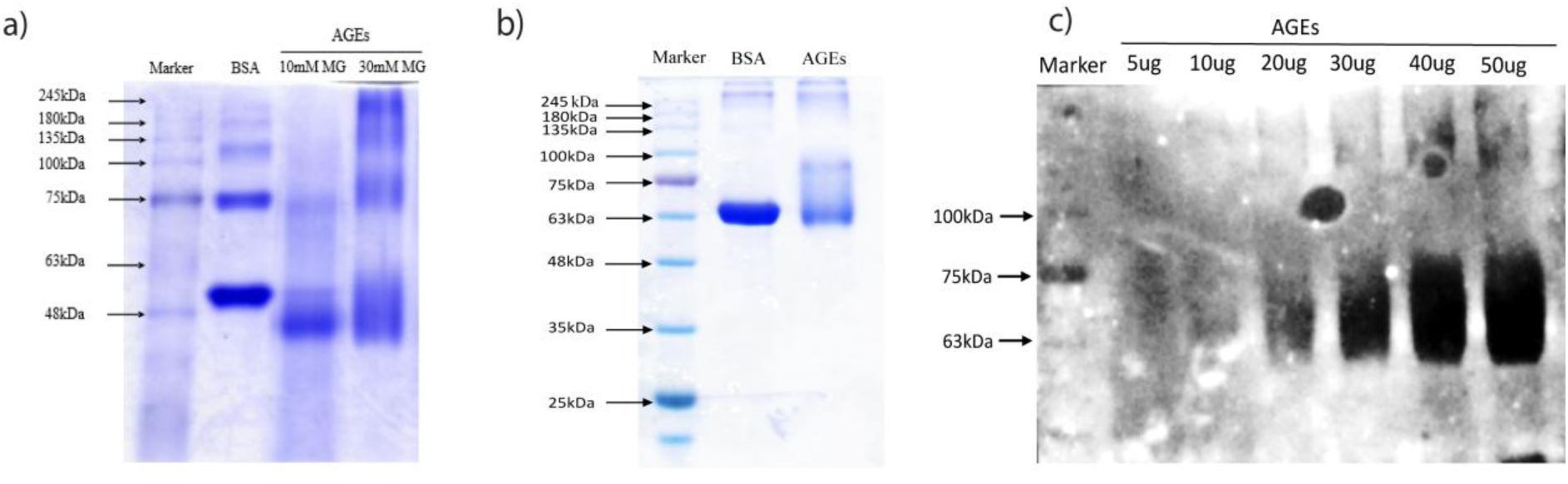
Characterization of MG-BSA (AGEs) by **(a)** CBB stained native PAGE, showing three bands in BSA control, and three diffused bands in AGEs. **(b)** CBB stained SDS-PAGE showing 66 kDa band in BSA control, mark the smear formation by AGEs of different molecular weights. **(c)** Western blot analysis of MG-BSA (AGEs) using Anti-AGEs antibody (ab23722). The thickness of bands increases with increasing concentration of AGEs (n=3).

### 3.6. Dynamic Light Scattering (DLS)

Dynamic light scattering (DLS) was used to study the aggregation induced in MG-BSA samples. DLS is a spectroscopic technique that measures the hydrodynamic radius of particles and makes it possible to measure the polydispersity of protein aggregates [37]. BSA has been reported to have a hydrodynamic radius of 3.5nm in its native form [38]. As shown in Figure 5a, BSA alone (SHAM) shows an increased hydrodynamic radius in the range of 5nm to 20nm (15.35 nm) with PDI 0.207. As temperature increases, BSA tends to form minor dimers, which are in equilibrium with monomers in solution, as a result of which a slight increase in polydispersity of sham BSA was also observed. Protein structure is dependent on many variables including solvent, pH, and temperature [36, 37]. Further interaction with MG leads to conformational changes in BSA, resulting in increased aggregation, the size of AGEs was observed to be 93.77 with PDI > 0.5 (Figure 5b), ranging from 100nm upto >300nm. Thus, BSA-MG complex shows polydispersed aggregates in a variable size range.

**Figure 5.**
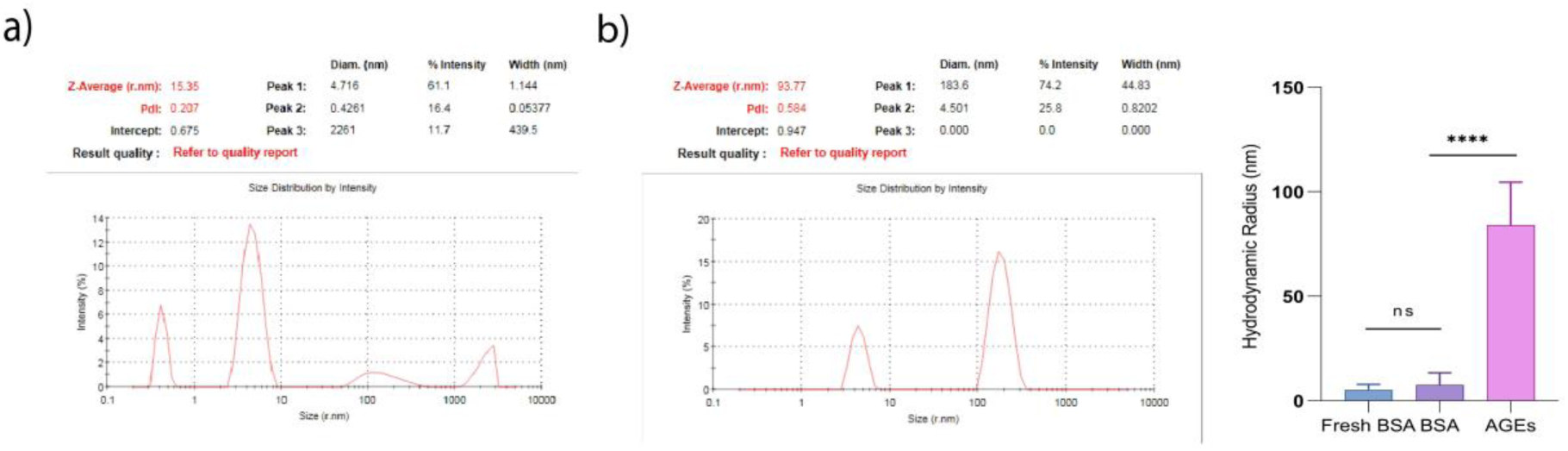
Dynamic light scattering size distribution curve of BSA (control) and MG-BSA (AGEs) obtained by Zeta Sizer (Malvern). BSA shows 15.35nm of hydrodynamic radius, whereas AGEs show an increase in radius with an average size of 93.77nm, Significance level ****p<0.0001, with respect to control (n=3).

### 3.7. Circular Dichroism

To investigate changes in secondary structural elements of MG-BSA (AGEs), circular dichroism (CD) spectroscopy was used. The technique is used to determine secondary structures, amyloid aggregates, and peptides [39]. For α-helical structures, circular dichroism (CD) spectra were characterised by two negative peaks in the far-UV region: one at approximately 208 nm and another at around 222 nm, and for β-sheets, a negative peak at approximately 218 nm is observed [40]. AGEs pool exhibited a mixture of helical and parallel sheet conformations, unlike native BSA, indicating that MG-induced glycation significantly alters structural integrity of BSA . MG-BSA (AGEs) shows an appreciable decrease in the α-helix (20.15 %) and an increase in the unordered structures and parallel sheets in the presence of 30 mM MG (Figure 6) [21].

**Figure 6.**
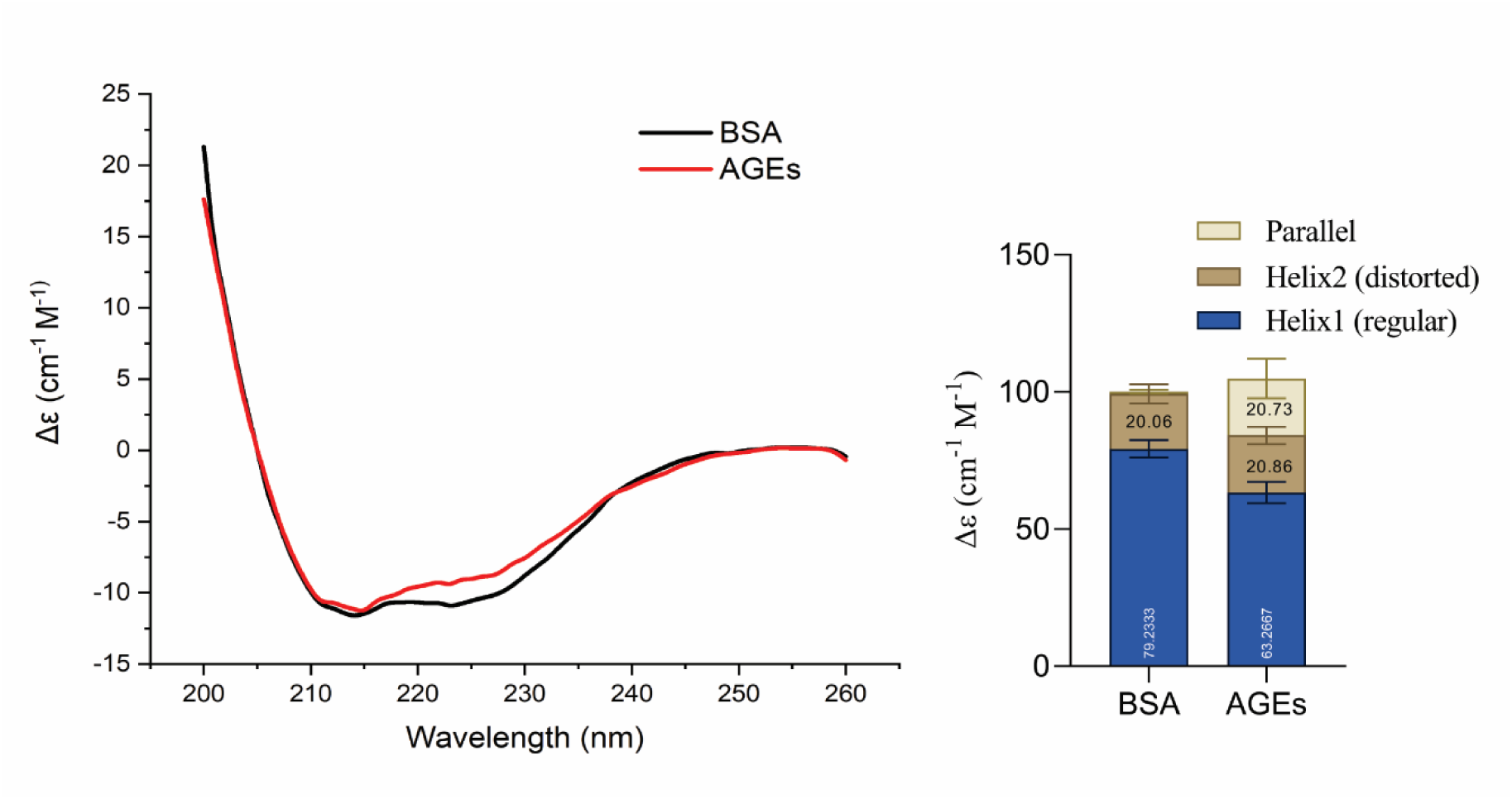
CD Spectrum of BSA and MG-BSA (AGEs) after incubation for 7 days. AGEs show a 20% increase in parallel structures (n=3).

### 3.8. Detection of aggregation using ThT and Congo-Red

ThT is a gold standard for amyloid detection and primarily binds along the shallow grooves on the beta-sheets, and fluorescence intensity increases with an increase in amyloid formation [41]. MG-BSA (AGEs) were found to exhibit increased fluorescence in comparison to control BSA (Figure 7a), suggesting conformational changes of BSA-AGEs. To validate the results further, the Congo red assay was also performed. Congo-red, a sulfonated hydrophobic molecule, binds to the exposed hydrophobic surfaces of partially folded proteins in β-pleated sheet conformation [42]. The increased absorbance at 490nm indicates structural changes in MG-BSA (AGEs), suggesting that CR is binding to aggregates (Figure 7b) [31]. Both CR assay and ThT assay results suggest structural/conformational changes in BSA due to MG.

**Figure 7.**
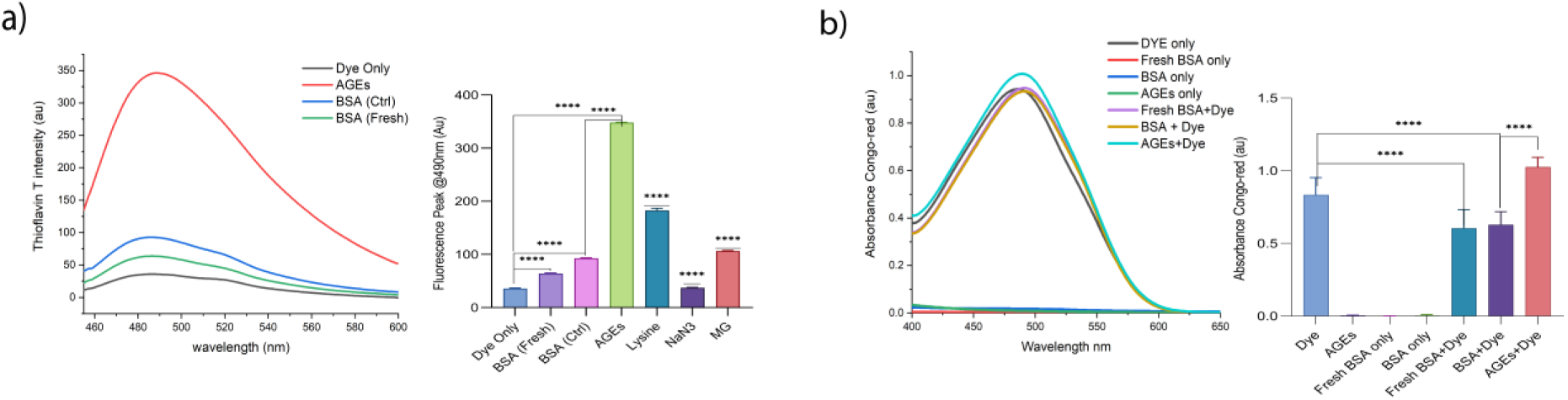
The effect of MG on BSA at 37°C for 7 days: **(a)** Increasing ThT fluorescence intensity of AGEs compared to native and control BSA. The spectrum was recorded at an excitation wavelength of 440nm, and emission was recorded at 490nm. **(b)** Congo-red (CR) UV-Vis absorbance, recorded within the range 400 nm-650 nm, Significance level ****p<0.0001, with respect to control (n=3).

### 3.9. Transmission Electron Microscopy and Atomic Force Microscopy

Transmission electron microscopy was performed for the morphological characterization of aggregates that are formed when BSA is incubated with MG for 7 days. TEM images of AGEs show dense oligomeric aggregates, which are absent in BSA control and fresh BSA (Figure 8a-c). This suggests that aggregation is induced by MG and is not due to self-aggregation of BSA. Largely, amyloid deposits are composed of bundles and fibres, but amorphous forms also exist [43–45].

**Figure 8.**
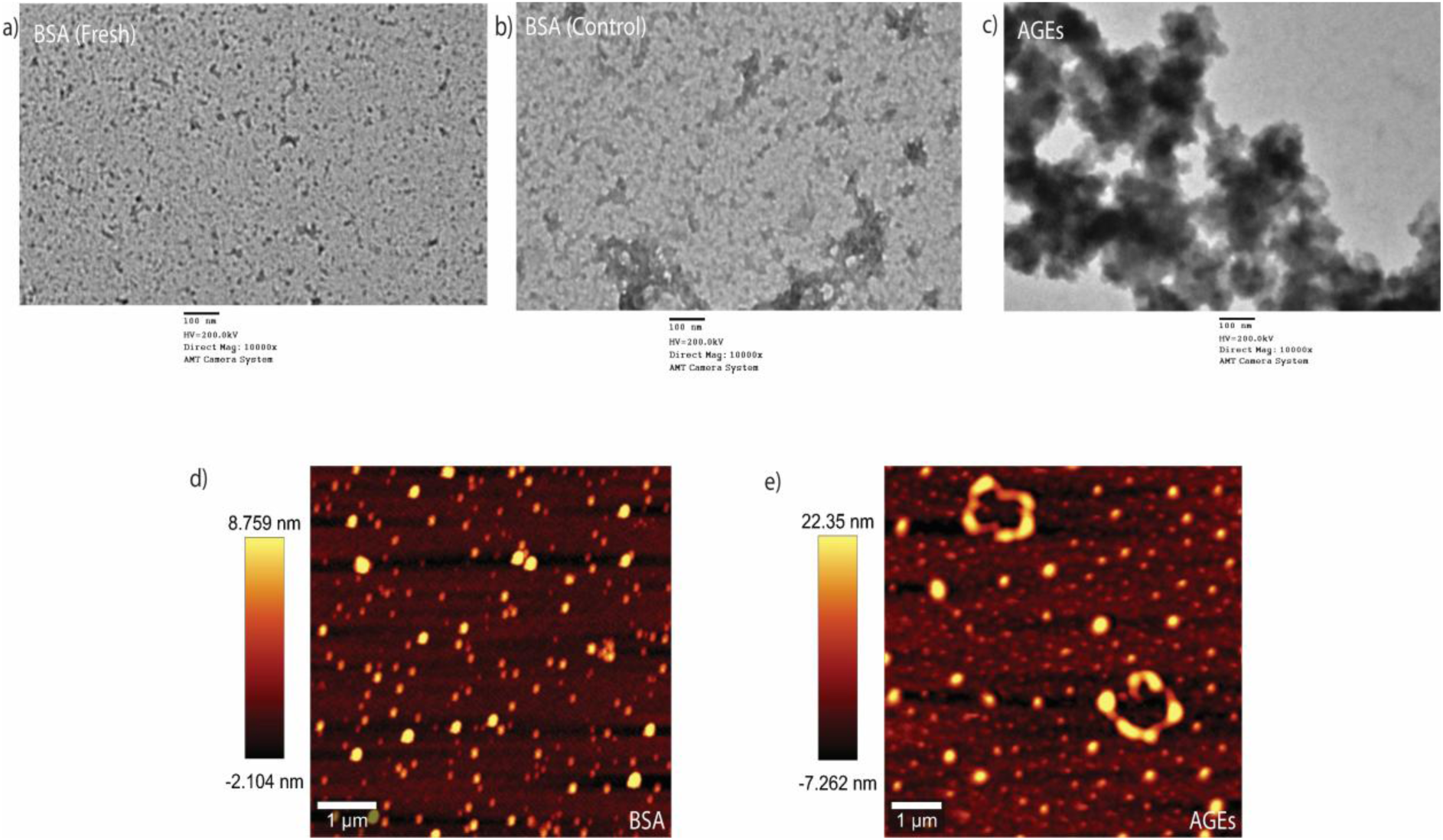
Effect of MG on BSA morphology: TEM images of **(a)** Fresh BSA, **(b)** BSA control, and **(c)** AGEs showing visible aggregation, AFM images of **(d)** BSA control, and **(e)** AGEs showing increased oligomers and height (n=3).

To further investigate the size of protein aggregates, AFM analysis was done and data showed BSA did not form a significant amount of aggregates and was in the size range of -2.104nm to 8.759 nm (Figure 8d). However, in MG-BSA spheroidal oligomers with increased height range, of -7.262nm to 22.35nm were observed (Figure 8e). These data suggest that MG leads to structural changes in protein (BSA), leading to the formation of higher-order oligomeric aggregates.

### 3.10. Effect of phytochemicals on MG-BSA (AGEs)

Advanced glycation end products (AGEs) not only correlate with age-related diseases and cancer, but their excessive accumulation also contributes to the pathogenesis of various other disease conditions. Therefore, we evaluated the effect of two phytochemicals, GA and CA, on AGEs formation *in vitro*. Following initial dose-response studies, the effect of phytochemicals (50 µM) was evaluated using fluorescence spectroscopy. Both the phytochemicals exhibited a decrease in AGEs formation as observed by a decrease in fluorescence (Figure 9a). Antiglycation abilities are often associated with antioxidant and anti-carbonyl properties [46]. GA, due to the greater number of hydroxyl groups on the flavonoid ring, enables stronger bonding interactions with radicals, serving as a better antioxidant/antiglycation agent than CA [47, 48]. A decrease in UV absorbance was also observed in GA and CA treated samples, validating fluorescence spectroscopy results (Figure 9b). ThT and CR staining confirmed the inhibitory effect on AGEs formation. (Figure 9c,d).

**Figure 9.**
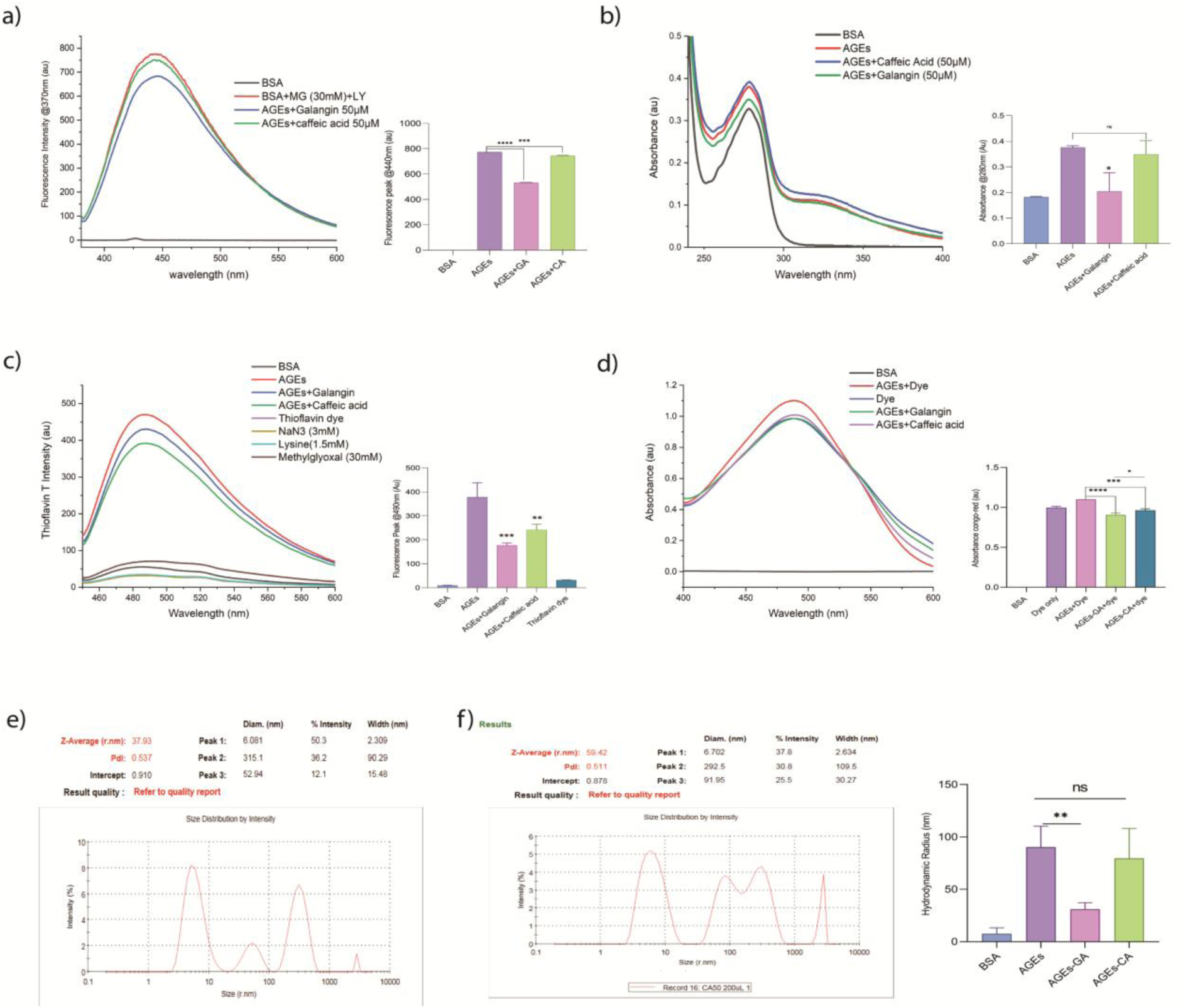
Antiglycation effect of GA and CA on AGEs formation: The fluorescence and UV-vis spectrum of AGEs, BSA control, and AGEs in the presence of **(a,b)** GA, and CA, respectively. Anti-aggregation effect of GA and CA using **(c)** ThT fluorescence spectrum, and **(d)** congo-red UV-vis spectrum. The dynamic light scattering size distribution curve of **(e)** GA and **(f)** CA was obtained by ZetaSizer (Malvern), Significance level ****p<0.0001, ***p<0.0005, **p<0.005, *p<0.05 with respect to control (n=3).

DLS analysis shows that the aggregates obtained after GA and CA treatment were significantly smaller in size than MG-BSA (AGEs). Treatment with GA shows a decrease in size to 37.9nm with a polydispersity index of 0.537, whereas CA shows a hydrodynamic radius of 59.42nm with PdI of 0.511 (Table 1; Figure 9e,f). This, strongly suggests that phytochemicals GA and CA have potent anti-glycation activity while Galangin is significantly more potent than Caffeic Acid.

**Table 1:** Effect of phytochemicals on AGEs formation.

| S.No. | Sample | Hydrodynamic Radius (r.nm) |
| --- | --- | --- |
| 1. | BSA | 5-20 |
| 2. | AGEs | >200 |
| 3. | AGEs-GA | 30-55 |
| 4. | AGEs-CA | 60-99 |

Thus, GA and CA show antiglycation as well as anti-aggregation properties. In our study we found GA as more potent antiglycation agent, even at lower concentrations in comparison to CA. Apart from antioxidative properties, CA is reported to mask Arg185 and Tyr137 residues using hydrogen bonds, prevent AGEs generation and also stabilise the structure of BSA [49] on the other hand, Galangin is reported to entrap MG and make Galangin-MG adducts [50]. It has also been reported that Galangin has the potential to protect free amino groups of the proteins [51]. The antiglycation activity of Galangin and Caffeic acid may also be due to free radical scavenging activity [50, 52]. In a study, it was also found that a phytochemical, α-Mangostin, not only inhibited AGEs formation but also suppressed β-sheet and β-turn components in α-glucosidase structure [53], indicating the potential of phytochemicals in Caffeic acid is reported to induce a reduction in α-glucosidase and α-amylase enzyme activity [54]. Caffeic acid and Galangin not only suppress AGEs formation but also inhibit fructosamine, which is an early glycation product [48]. The inhibition of Galangin on fructosamine formation can be explained by its metal ion chelation, which inhibits Amadori product formation [55].

## Conclusion

AGEs were formed *in vitro* by incubating BSA with methylglyoxal. The formation of BSA-MG aggregates was confirmed by fluorescence spectroscopy, UV Vis spectroscopy and Circular Dichroism. Formation of amyloid-like aggregates was further confirmed by Thioflavin T and Congo Red assays. These *in vitro* AGEs were used to check the antiglycation potential of phytochemicals, GA and CA, that have already been reported to have anticancer properties. They were also found to show antiglycation activity in the present study. Their antiglycating effect was confirmed using various parameters. A decrease in protein aggregation and AGEs formation was observed. This could be a successful strategy to evaluate phytochemicals for protein aggregation-based diseases like neurodegenerative disorders, where protein deposits are central to pathology, as well as cancers, where, due to the Warburg effect, excessive AGEs are produced, promoting tumour growth and invasiveness.

## Data Availability

Data will be made available on request.

## Funding Received

This work was supported by grant from Indian Council of Medical Research to A.B. Tiku (36/10/2020-Toxi/BMS) and CSIR fellowship (09/263(1225)/2020-EMR-I) to Nupur Kanojia. We acknowledge the facilities/laboratories provided by DBT BUILDER (grant no. BT/INF/22/SP45382/2022) and DST FIST-II [grant no. SR/FST/LSII-046/2016(C)].

## Authors and Affiliations

**Nupur Kanojia and A.B. Tiku**

Radiation and Cancer Therapeutics Laboratory, School of Life Science, Jawaharlal Nehru University, New Delhi,110067, India.

## Contribution

ABT: Conceptualisation and Funding Acquisition, Review and manuscript writing.

NK: Performed experiments, analysed data, drafted manuscript,

Both authors reviewed the manuscript.

## Corresponding author

A.B. Tiku

## Ethical Declarations

The authors declare no competing interests

## References

[1] N. Rabbani, P.J.J.R.B. Thornalley, Protein glycation–biomarkers of metabolic dysfunction and early-stage decline in health in the era of precision medicine, 42 (2021) 101920.

[2] K. Waqas, M. Muller, M. Koedam, Y. El Kadi, M.C. Zillikens, B.J.B. Van der Eerden, Methylglyoxal–an advanced glycation end products (AGEs) precursor–Inhibits differentiation of human MSC-derived osteoblasts in vitro independently of receptor for AGEs (RAGE), 164 (2022) 116526.

[3] C. Nigro, A. Nicolò, I. Prevenzano, P. Formisano, F. Beguinot, C.J.O. Miele, The dual-role of methylglyoxal in tumor progression—Novel therapeutic approaches. Front, 11(645686) (2021) 10.3389.

[4] S. Yamagishi, K. Nakamura, H.J.M.h. Inoue, Therapeutic potentials of unicellular green alga Chlorella in advanced glycation end product (AGE)-related disorders, 65(5) (2005) 953–955.

[5] O.O. Omofuma, D.P. Turner, L.L. Peterson, A.T. Merchant, J. Zhang, S.E.J.C.P.R. Steck, Dietary advanced glycation end-products (AGE) and risk of breast cancer in the prostate, lung, colorectal and ovarian cancer screening trial (PLCO), 13(7) (2020) 601–610.

[6] A.J.B.R. Kuzan, Toxicity of advanced glycation end products, 14(5) (2021) 46.

[7] D. Schröter, A.J.C.p.d. Höhn, Role of advanced glycation end products in carcinogenesis and their therapeutic implications, 24(44) (2018) 5245–5251.

[8] B. Dariya, G.P.J.D.D.T. Nagaraju, Advanced glycation end products in diabetes, cancer and phytochemical therapy, 25(9) (2020) 1614–1623.

[9] Y. Zhang, Z. Zhang, C. Tu, X. Chen, R.J.A. He, Advanced glycation end products in disease development and potential interventions, 14(4) (2025) 492.

[10] D. Tagliazucchi, S. Martini, A.J.J.o.A. Conte, F. Chemistry, Protocatechuic and 3, 4-dihydroxyphenylacetic acids inhibit protein glycation by binding lysine through a metal-catalyzed oxidative mechanism, 67(28) (2019) 7821–7831.

[11] Deepika, P.K.J.B. Maurya, Ellagic acid: insight into its protective effects in age-associated disorders, 12(12) (2022) 340.

[12] A. Ouamnina, A. Alahyane, I. Elateri, M. Ouhammou, M.J.H. Abderrazik, In vitro and molecular docking studies of antiglycation potential of phenolic compounds in date palm (Phoenix dactylifera L.) fruit: exploring local varieties in the food industry, 10(6) (2024) 657.

[13] A. Febriza, A.A. Zahrah, N.S. Andini, F. Usman, H.H.J.D. Idrus, Metabolic Syndrome, Obesity, Potential Effect of Curcumin in Lowering Blood Glucose Level in Streptozotocin-Induced Diabetic Rats, (2024) 3305–3313.

[14] Z. Wei, Q.J.B.J.S.T.R. Huang, Adverse health consequences of dietary advanced glycation end products (AGEs) and inhibitory effects of natural ingredients on ages, 1 (2017) 1386–90.

[15] U.J. Jung, M.-K. Lee, Y.B. Park, S.-M. Jeon, M.-S.J.J.o.p. Choi, e. therapeutics, Antihyperglycemic and antioxidant properties of caffeic acid in db/db mice, 318(2) (2006) 476–483.

[16] H.S. Tuli, K. Sak, S. Adhikary, G. Kaur, D. Aggarwal, J. Kaur, M. Kumar, N.C. Parashar, G. Parashar, U.J.E.B. Sharma, Medicine, Galangin: A metabolite that suppresses anti-neoplastic activities through modulation of oncogenic targets, 247(4) (2022) 345–359.

[17] M.S. Rahaman, M.A. Siraj, M.A. Islam, P.C. Shanto, O. Islam, M.A. Islam, J.J.T.J.o.N.B. Simal-Gandara, Crosstalk between xanthine oxidase (XO) inhibiting and cancer chemotherapeutic properties of comestible flavonoids-a comprehensive update, 110 (2022) 109147.

[18] K.M.M. Espíndola, R.G. Ferreira, L.E.M. Narvaez, A.C.R. Silva Rosario, A.H.M. Da Silva, A.G.B. Silva, A.P.O. Vieira, M.C.J.F.i.o. Monteiro, Chemical and pharmacological aspects of caffeic acid and its activity in hepatocarcinoma, 9 (2019) 541.

[19] S. Celik, S. Erdogan, M.J.P.R. Tuzcu, Caffeic acid phenethyl ester (CAPE) exhibits significant potential as an antidiabetic and liver-protective agent in streptozotocin-induced diabetic rats, 60(4) (2009) 270–276.

[20] A. Bhatwadekar, V.J.J.o.c.l.a. Ghole, Rapid method for the preparation of an AGE-BSA standard calibrator using thermal glycation, 19(1) (2005) 11–15.

[21] A. Ahmed, A. Shamsi, M.S. Khan, F.M. Husain, B.J.I.j.o.b.m. Bano, Methylglyoxal induced glycation and aggregation of human serum albumin: Biochemical and biophysical approach, 113 (2018) 269–276.

[22] R.L. Levine, D. Garland, C.N. Oliver, A. Amici, I. Climent, A.-G. Lenz, B.-W. Ahn, S. Shaltiel, E.R. Stadtman, [49] Determination of carbonyl content in oxidatively modified proteins, Methods in enzymology, Elsevier 1990, pp. 464–478.

[23] U.K.J.n. Laemmli, Cleavage of structural proteins during the assembly of the head of bacteriophage T4, 227(5259) (1970) 680–685.

[24] K.P. Prajapati, A.P. Singh, K. Dubey, M. Ansari, M. Temgire, B.G. Anand, K.J.C. Kar, S.B. Biointerfaces, Myricetin inhibits amyloid fibril formation of globular proteins by stabilizing the native structures, 186 (2020) 110640.

[25] M. Mesías, C.J.C.O.i.F.S. Delgado-Andrade, Melanoidins as a potential functional food ingredient, 14 (2017) 37–42.

[26] I. Berdowska, M. Matusiewicz, I.J.M. Fecka, Methylglyoxal in Cardiometabolic disorders: Routes leading to pathology counterbalanced by treatment strategies, 28(23) (2023) 7742.

[27] A. Twarda-Clapa, A. Olczak, A.M. Bia³kowska, M.J.C. Kozio³kiewicz, Advanced glycation end-products (AGEs): Formation, chemistry, classification, receptors, and diseases related to AGEs, 11(8) (2022) 1312.

[28] H.J.M. Kataoka, Current Developments in Analytical Methods for Advanced Glycation End Products in Foods, 30(20) (2025) 4095.

[29] J. Ahmad, K.J.R.A. Alam, Impact of in vitro non-enzymatic glycation on biophysical and biochemical regimes of human serum albumin: relevance in diabetes associated complications, 5(78) (2015) 63605–63614.

[30] A. Schmitt, J. Schmitt, G. Münch, J.J.A.b. Gasic-Milencovic, Characterization of advanced glycation end products for biochemical studies: side chain modifications and fluorescence characteristics, 338(2) (2005) 201–215.

[31] I. Dalle-Donne, R. Rossi, D. Giustarini, A. Milzani, R.J.C.c.a. Colombo, Protein carbonyl groups as biomarkers of oxidative stress, 329(1-2) (2003) 23–38.

[32] R. Nagai, K. Matsumoto, X. Ling, H. Suzuki, T. Araki, S.J.D. Horiuchi, Glycolaldehyde, a reactive intermediate for advanced glycation end products, plays an important role in the generation of an active ligand for the macrophage scavenger receptor, 49(10) (2000) 1714–1723.

[33] A.R. Mir, K. Alam, A.J.I.j.o.b.m. Ali, Methylglyoxal mediated conformational changes in histone H2A—generation of carboxyethylated advanced glycation end products, 69 (2014) 260–266.

[34] J.-H.J.B.r. Kang, Oxidative modification of human ceruloplasmin by methylglyoxal: an in vitro study, 39(3) (2006) 335–338.

[35] C. Li, T.J.I.j.o.b.m. Arakawa, Application of native polyacrylamide gel electrophoresis for protein analysis: Bovine serum albumin as a model protein, 125 (2019) 566–571.

[36] J.T. Wu, M.C. Tu, P.J.J.o.c.l.a. Zhung, Advanced glycation end product (AGE): Characterization of the products from the reaction between D-glucose and serum albumin, 10(1) (1996) 21–34.

[37] Y. Li, G. Yang, Z.J.A.P.S.B. Mei, Spectroscopic and dynamic light scattering studies of the interaction between pterodontic acid and bovine serum albumin, 2(1) (2012) 53–59.

[38] S. Falke, K. Dierks, C. Blanchet, M. Graewert, F. Cipriani, R. Meijers, D. Svergun, C.J.J.o.s.r. Betzel, Multi-channel in situ dynamic light scattering instrumentation enhancing biological small-angle X-ray scattering experiments at the PETRA III beamline P12, 25(2) (2018) 361–372.

[39] W.D. McCubbin, C. Kay, S. Narindrasorasak, R.J.B.J. Kisilevsky, Circular-dichroism studies on two murine serum amyloid A proteins, 256(3) (1988) 775–783.

[40] G.R. Upchurch, N. Ramdev, M.T. Walsh, J.J.J.o.t. Loscalzo, thrombolysis, Prothrombotic consequences of the oxidation of fibrinogen and their inhibition by aspirin, 5 (1998) 9–14.

[41] C. Wu, M. Biancalana, S. Koide, J.-E.J.J.o.m.b. Shea, Binding modes of thioflavin-T to the single-layer â-sheet of the peptide self-assembly mimics, 394(4) (2009) 627–633.

[42] R. Khurana, V.N. Uversky, L. Nielsen, A.L.J.J.o.B.C. Fink, Is Congo red an amyloid-specific dye?, 276(25) (2001) 22715–22721.

[43] E.I. Yakupova, L.G. Bobyleva, I.M. Vikhlyantsev, A.G.J.B.r. Bobylev, Congo Red and amyloids: history and relationship, 39(1) (2019) BSR20181415.

[44] J.D. Sipe, A.S.J.J.o.s.b. Cohen, History of the amyloid fibril, 130(2-3) (2000) 88–98.

[45] G.S.J.A.i.a.p. Markowitz, Dysproteinemia and the kidney, 11(1) (2004) 49–63.

[46] W. Wang, Y. Yagiz, T.J. Buran, C. do Nascimento Nunes, L.J.F.r.i. Gu, Phytochemicals from berries and grapes inhibited the formation of advanced glycation end-products by scavenging reactive carbonyls, 44(9) (2011) 2666–2673.

[47] F. Caruso, M. Berinato, M. Hernandez, S. Belli, C. Smart, M.J.P.O. Rossi, Antioxidant properties of bee propolis and an important component, galangin, described by X-ray crystal structure, DFT-D and hydrodynamic voltammetry, 17(5) (2022) e0267624.

[48] M.S. Khan, M.S. Alokail, A.M.H. Alenad, N. Altwaijry, N.O. Alafaleq, A.M. Alamri, M.A.J.M. Zawba, Binding studies of caffeic and p-coumaric acid with α-amylase: Multispectroscopic and computational approaches deciphering the effect on advanced glycation end products (AGEs), 27(13) (2022) 3992.

[49] X. Cao, Y. Xia, M. Zeng, W. Wang, Y. He, J.J.C. Liu, biodiversity, Caffeic acid inhibits the formation of advanced glycation end products (AGEs) and mitigates the AGEs-induced oxidative stress and inflammation reaction in human umbilical vein endothelial cells (HUVECs), 16(10) (2019) e1900174.

[50] Z. Sheng, B. Ai, L. Zheng, X. Zheng, Z. Xu, Y. Shen, Z.J.I.J.o.F.S. Jin, Technology, Inhibitory activities of kaempferol, galangin, carnosic acid and polydatin against glycation and á-amylase and á-glucosidase enzymes, 53(3) (2018) 755–766.

[51] X. Peng, K.-W. Cheng, J. Ma, B. Chen, C.-T. Ho, C. Lo, F. Chen, M.J.J.o.a. Wang, f. chemistry, Cinnamon bark proanthocyanidins as reactive carbonyl scavengers to prevent the formation of advanced glycation endproducts, 56(6) (2008) 1907–1911.

[52] S. Son, B.A.J.J.o.a. Lewis, f. chemistry, Free radical scavenging and antioxidative activity of caffeic acid amide and ester analogues: Structure− activity relationship, 50(3) (2002) 468–472.

[53] F.M. Djeujo, V. Francesconi, M. Gonella, E. Ragazzi, M. Tonelli, G.J.M. Froldi, Anti-α-glucosidase and antiglycation activities of α-mangostin and new xanthenone derivatives: enzymatic kinetics and mechanistic insights through in vitro studies, 27(2) (2022) 547.

[54] K. Kannan, S.J.R.a. Sadhukhan, A multimodal caffeic acid-derived alkyl-amide antidiabetic agent: targeting á-glucosidase, á-amylase, oxidative stress, and protein glycation, 15(49) (2025) 41833–41849.

[55] L. Zeng, H. Ding, X. Hu, G. Zhang, D.J.F.C. Gong, Galangin inhibits á-glucosidase activity and formation of non-enzymatic glycation products, 271 (2019) 70–79.

